# Assessing bactericidal dynamics of persister-manifested clinical isolates in the presence of macrophages and antibiotics

**DOI:** 10.64898/2025.12.22.696100

**Authors:** Jia Hao Yeo, Nasren Begam, Wan-Ting Leow, Yang Zhong, Andrea Lay-Hoon Kwa

**Author notes:** **Correspondence** to Andrea Lay-Hoon Kwa,.

## Abstract

Antibiotics are often depicted to reduce bacterial load for immuno-clearance. Recurring infections can be caused by bacteria persisters that survived antibiotic treatment. It is unclear how the host immunity manages infection by persister-manifested strains during antibiotic treatment. Differentiated THP-1 macrophages were exposed to persister-manifested clinical isolates at various multiplicities-of-infection (MOI), in the presence or absence of meropenem. Across all MOIs, meropenem did not greatly reduce the intracellular bacterial load nor eradicate all extra-macrophagic non-persister bacteria. Bactericidal effects on extracellular bacteria vary for each strain after meropenem treatment in mildly infected macrophages. In severely infected macrophages, extra-macrophagic bacteria of all isolates decreases in numbers followed by a bacteriostatic-like effect. Our findings contradict the common notion that antibiotics generally reduce the bacterial load for immune clearance. Our study suggests that the bactericidal effects of meropenem differs with host presence and can be dependent on bacterial load.

## 1. INTRODUCTION

Antibiotics are widely recognized to reduce overall bacterial load but often fail to achieve complete bacterial eradication *in vivo*.[1–4] Standard antibiotics efficacy is typically evaluated based on the minimum inhibitory concentrations (MICs) using bacteria isolates at a fixed inoculum *in vitro*, without considering the host. It remains unclear if antibiotics exhibit altered efficacy across a spectrum of different bacterial loads, in the presence of host. Bacteria load, bacterial physiology, and the host environment are likely interrelated determinants of antibiotics efficacy.

During an infection, the bacterial load influences both the severity of disease and the effectiveness of host immune responses.[5,6] The bacterial load at the onset of infection (also known as the inoculum size) can dictate the progression of the infection. A high bacterial load can exacerbate immune activation and overwhelm host defences. Conversely, infections initiated by a low bacterial load can still progress, possibly due to mechanisms such as immune evasion or high bacterial virulence.[7–12] As infection progresses, the bacterial load contributes to the overall bacterial burden, which can vary across individuals and tissues, and affected by the presence of immune or pharmacologic pressures such as antibiotics.

The complexity of host–pathogen interactions increases further when the pathogen is manifested with heterogeneous populations, such as presence of bacteria persisters. Persisters are a sub-population of antibiotic susceptible bacteria that survive antibiotics at bactericidal concentrations, without acquiring genetic resistance.[13–16] These persisters can be resuscitated after antibiotic removal and contribute to recurrent or persistent bacterial infections.[17,18] Recently, there are increasing studies examining host-bacterial persisters interactions of various microbial species.[19–27] However, it is still unclear how host immunity manages infections caused by persister-containing bacterial populations during antibiotic treatment. This is analogous to a human patient receiving antibiotic treatments while still suffering from a persistent bacterial infection. Growth resuscitation of persisters upon antibiotic removal can cause recurrent infections, which are common clinically. As such, we want to find out if an immunocompetent patient has the ability to eradicate persister emerging from antibiotic treatment.

The formation and abundance of persister cells vary depending on the antibiotic used, even within the same bacterial strain.[28] Different antibiotics can induce persister cells through distinct physiological pathways.[28,29] In *Escherichia coli*, a study showed no consistent correlation in persister levels across antibiotics, even those with similar modes of action like ciprofloxacin and nalidixic acid.[30]

In our previous studies, screening of *E.* coli clinical isolates revealed persister cells phenotypes following meropenem treatment.[31] These persisters were identified using a set of criteria based on the consensus established made by multiple leading research groups.[14,31] Following this study that examined only with meropenem, we aim to assess the bactericidal dynamics of persister-manifested clinical isolates when treated with antibiotics, in the presence of the immune cells. We hypothesized that meropenem could both greatly reduce the bacterial load within macrophages and eradicate all non-persister bacteria.

## 2. MATERIALS AND METHODS

### 2.1 Clinical isolates

Nonclonal clinical strains of *E. coli* were previously collected as part of a surveillance study from 2011 to 2018 (originally obtained from Singapore General Hospital’s Diagnostic Bacteriology Laboratory, Singapore). Bacterial strains were stored at -80°C in Cryobank (Thermo Scientific, Singapore) storage vials. Fresh isolates were sub-cultured twice on 5% sheep blood agar plates (Thermo Scientific, Singapore) for 24 h at 35°C before each experiment.

### 2.2 Cell line and maintenance

Human THP-1 cells were obtained as a gift from Prof. Linfa Wang (Duke-NUS, Singapore) and previously acquired from American Type Culture Collection (ATCC TIB-202). The promonocytic THP-1 cell line, was cultured in RPMI-1640 media (Life Technologies, Grand Island, NY, USA) supplemented only with 10% fetal bovine serum (Life Technologies, Gibco, Grand Island, NY, USA). THP-1 cells were maintained at logarithmic growth phase in a 37^0^C incubator supplemented with 5% CO_2_. Cells were passaged twice weekly at a cell density of 5 x 10^5^ cells in a T75 cm^2^ flask.

### 2.3 Differentiation of THP-1 cells to macrophages

Prior to the differentiation, THP-1 cells were sub-cultured in a T175 cm^2^ flask at a cell density of 1 x 10^6^ cells to obtain sufficient cells for the macrophage infection assay performed in triplicates. THP-1 cells were then seeded at a cell density of 1 x 10^5^ in each well of a 6-well culture dish. The promonocytic THP-1 cells were then differentiated into macrophages in differentiation media (RPMI-1640 media containing 50 ng/mL phorbol-12-myristate-13-acetate) over 3 days [32].

### 2.4 Macrophage infection assay

Differentiated THP-1 macrophages seeded in 6-well culture dishes were exposed to clinical isolate at various multiplicities of infections (MOIs; bacteria-to-macrophage ratio) in triplicates. Culture dishes containing macrophages with the infecting bacteria were then placed in a shaking incubator at 35^0^C incubator. At each timepoint, both extra-macrophagic (extracellular) and intra-macrophagic (intracellular) bacteria were harvested and colonies were quantified via viable plating. After harvesting extracellular bacteria, adhered macrophages were gently washed with phosphate buffered saline five times. Each well of the culture dish was then checked visually using an inverted microscope over multiple fields of view to ensure only macrophages remained. Intracellular bacteria were next harvested by permeabilising macrophages with 0.1% (v/v) Triton-X-100 [33], followed by mechanical lysing using cell scrappers. Each well of the culture dish was checked visually again using an inverted microscope over multiple fields of view to ensure no macrophage remains.

### 2.5 Macrophage infection assay with antibiotics

Assay was performed with modifications to the protocol described above. Briefly, culture dishes containing the macrophages with the infecting bacteria were placed in a shaking incubator at 35^0^C incubator for 2 hours to reach saturation. In simulating antibiotic treatment, meropenem (antibiotic; Toronto Research Centre, Toronto, Canada) was then added at a clinical achievable concentration of 20 mg/L [31,34]. At each timepoint after adding meropenem, both extra-macrophagic (extracellular) and intra-macrophagic (intracellular) bacteria were harvested and colonies were quantified via viable plating.

### 2.6 Viable plating and colony enumeration

Harvested bacteria (100 μL) were spirally plated onto Müeller-Hinton agar plates (Thermo Scientific, Singapore).). Agar plates were incubated overnight at 35°C for at least 18 h to 24 h. Colonies formed were enumerated visually. The lower limit of detection for the viable colony counts was 2.4 log_10_ CFU/mL.

### 2.7 Lactate dehydrogenase release assay

Lactate dehydrogenase release assay was performed according to protocol described by the manufacturer (Roche, Basel, Switzerland) in the Cytotoxicity Detection Kit. Absorbance of the resultant formazan product derived from the assay was measured using the EnSpire^TM^ Spectrometer (PerkinElmer, Waltham, MA, USA) complemented with the acquiring software, EnSpire Workstation (version 4.13.3005.1482).

### 2.8 Determining minimum inhibitory concentrations

A standard broth dilution method was used to determine MIC of meropenem for the isolates used in this study. All assessments were performed under CLSI guidelines.[35]

### 2.9 Statistical analyses

An unpaired two-tailed *t*-test was performed to compare differences between two groups. Analyses were conducted using GraphPad QuickCalcs (GraphPad Software, La Jolla, CA; https://www.graphpad.com/quickcalcs/ttest1/, accessed on May 2025). A *P*-value < 0.05 was considered statistically significant.

## 3. RESULTS

### 3.1 Determining the duration in reaching bacterial saturation load in THP-1 macrophages

Human THP-1 cells were differentiated into macrophages using 50 ng/mL phorbol-12-myristate-13-acetate over 3 days [32]. Differentiated THP-1 macrophages were exposed to each clinical isolate at various multiplicities-of-infections (MOIs; bacteria-to-macrophage ratio. At each timepoint, both extracellular and intracellular bacteria were harvested and quantified using viable plating. Intracellular bacteria were harvested by permeabilising macrophages with 0.1% (v/v) Triton-X-100, followed by mechanical lysing using cell scrappers.

Bacterial uptake by macrophages were observable across all isolates after 1-hour post-infection at all MOIs (**Figure-1, Figure S1**). Viable intracellular bacteria numbers in macrophages reveal hyperbolic trends that did not suggest bactericidal event. Plateaux in viable bacteria counts in macrophages were consistently observed between 1-3 hours post infection. Therefore, we deemed 2-hours as the duration necessary to reach bacterial load saturation in THP-1 macrophages. The duration to reach plateaux is independent of the bacterial load.

**Figure 1:**
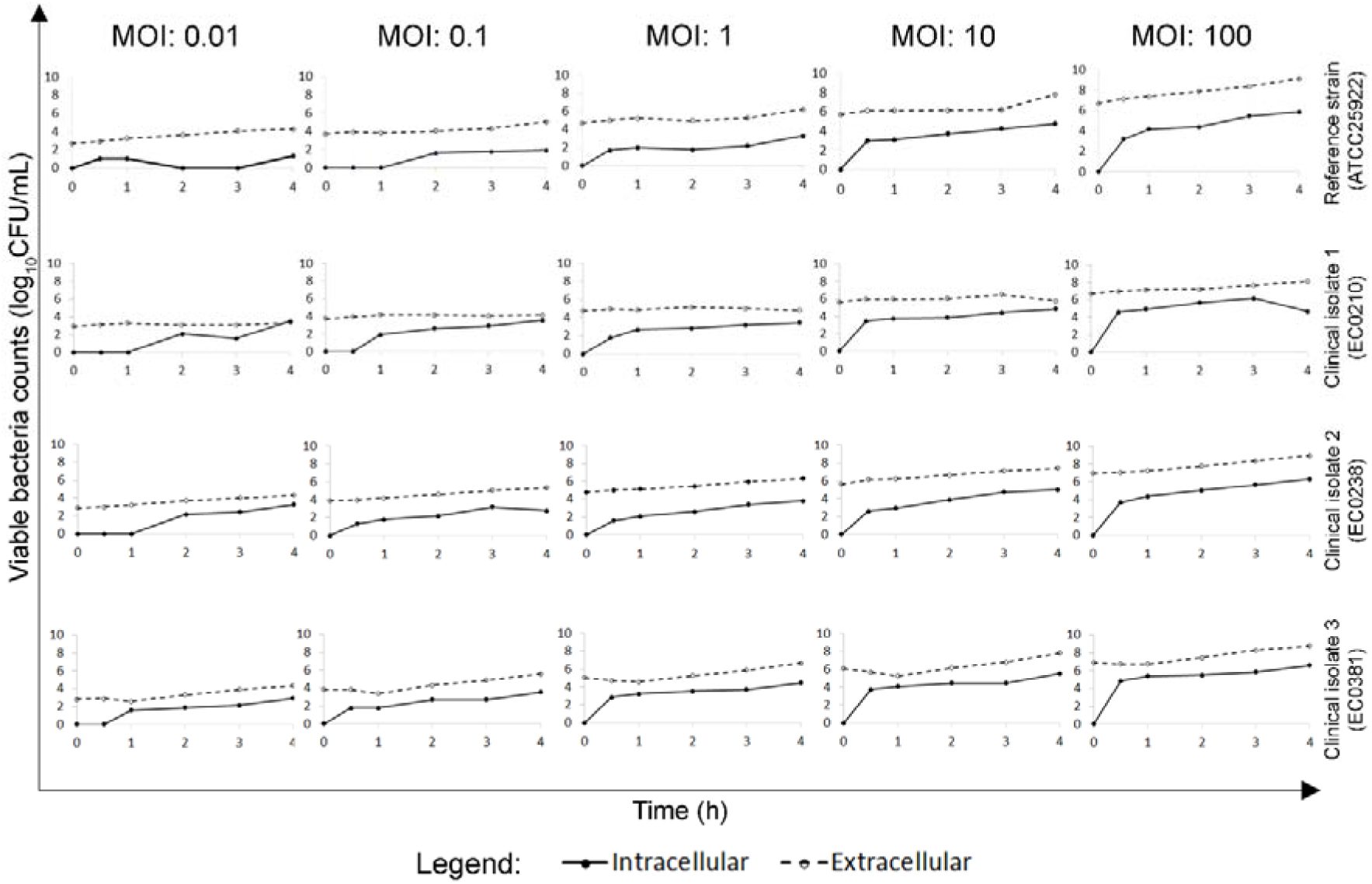
Bacteria load saturated in THP-1 macrophages at 2 hours. Differentiated THP-1 macrophages were exposed to different bacteria strains at varying multiplicity of infections (MOIs). MOIs refer to the bacteria-to-macrophage ratio. The highest MOI=100 refers to exposing 100-bacteria-to-1-macrophage, whilst the lowest MOI=0.01 refers to exposing 1-bacteria-to-100-macrophages. The ATCC25922 *Escherichia coli* was used as a reference strain in optimising the macrophage infection assay. At every time point, extracellular bacteria were harvested first. The adhered infected macrophages were then washed 5 times to ensure no remanent visible extracellular bacteria, as validated by visualising using an inverted microscope. Infected macrophages were then permeabilised with 0.1% (v/v) Triton-X-100 and lysed mechanically using a cell culture scraper. Harvested extracellular or intracellular bacteria were plated on Müeller Hinton agar plates. Colonies formed were enumerated the next day and graphed.

### 3.2 Bacteriostatic-like effects observed with meropenem treatment

Persisters are a sub-population of bacteria that becomes dormant upon antibiotic stress and can survive appropriate antibiotic treatment [14]. Growth resuscitation of persisters upon antibiotic removal can cause recurrent infections, which are common in clinic. As such, we want to find out if an immunocompetent patient has the ability to eradicate persister emerging from antibiotic treatment.

To simulate antibiotic treatment against bacterial infection *in-vitro*, macrophages were first infected for 2-hours to saturation load (**Figure-2A**). Thereafter, meropenem was added at a final, clinically achievable concentration (20 mg/L).[34] Lactate dehydrogenase release assay conducted reveal uninfected THP-1 macrophages were still viable (>90%) at 24-hour post-meropenem treatment. The meropenem MIC for these clinical isolates falls within the range of 4-8 mg/L, below 20 mg/L meropenem (≥ 2.5 x MIC) (**Table 1**) [31]. While these isolates are non-susceptible to meropenem, a higher dose of meropenem can still be used to eradicate the bacteria [36,37], in immunocompetent patients.

**Figure 2:**
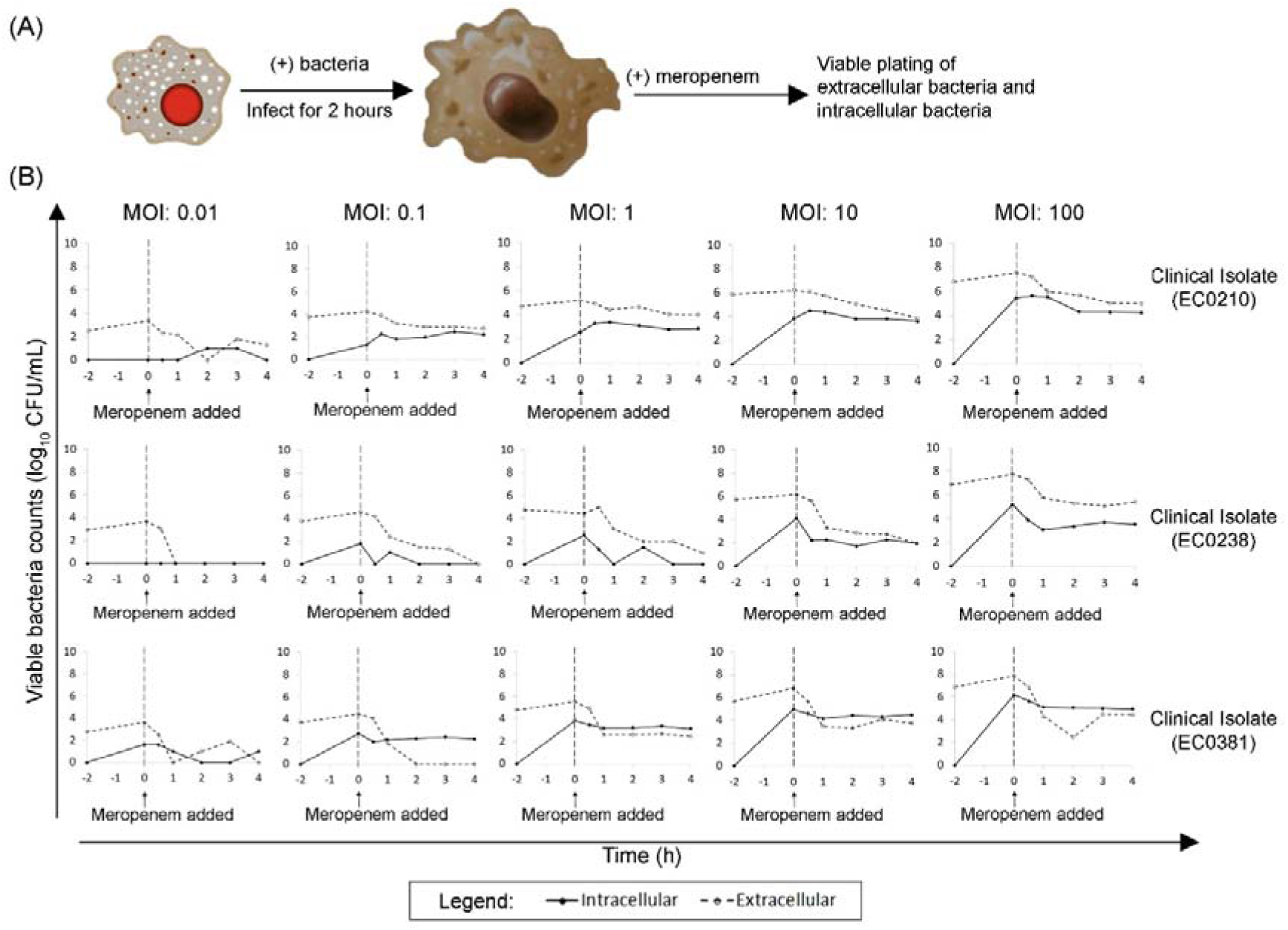
Bacteriostatic-like effects observed after 1-hour of meropenem treatment. (A) Schematic diagram of antibiotic treatment to an infected macrophage model is shown. (B) Differentiated THP-1 macrophages were exposed to different bacteria strains at varying multiplicity of infections (MOIs). MOIs refer to the bacteria-to-macrophage ratio. The highest MOI=100 refers to exposing 100-bacteria-to-1-macrophage, whilst the lowest MOI=0.01 refers to exposing 1-bacteria-to-100-macrophages. Macrophages were loaded with an clinical isolate till saturation for 2 hours, before adding meropenem to a final concentration of 20 mg/L. At every time point, extracellular bacteria were harvested first. The adhered infected macrophages were then washed 5 times to ensure no remanent visible extracellular bacteria, by visualising using an inverted microscope. Infected macrophages were then permeabilised with 0.1% (v/v) Triton-X-100 and lysed mechanically using a cell culture scraper. Harvested extracellular or intracellular bacteria were plated on Müeller Hinton agar plates. Colonies formed were enumerated the next day and graphed. Dotted line represents addition of meropenem after 2 hours post-infection, at bacterial saturation load.

**Table 1:**
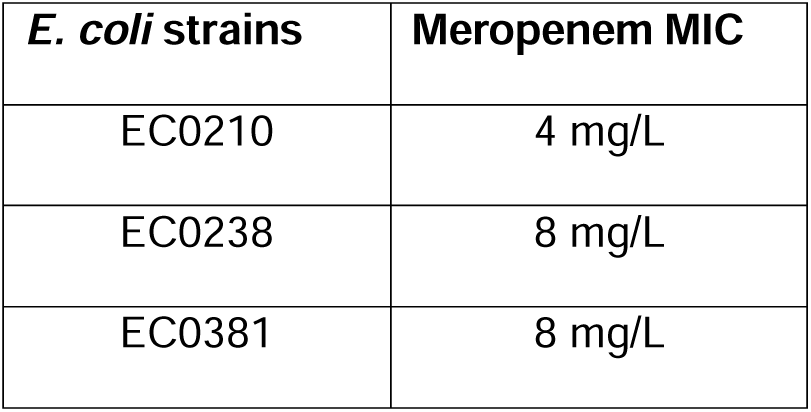
Meropenem MICs for each of the *E. coli* clinical isolates used. These clinical isolates were previous shown to harbour persisters upon meropenem treatment.

| *E. coli* strains | Meropenem MIC |
| --- | --- |
| EC0210 | 4 mg/L |
| EC0238 | 8 mg/L |
| EC0381 | 8 mg/L |

At various timepoints, both extracellular and intracellular bacteria were harvested and quantified using viable plating (**Figure-2B**). Against our hypothesis, across all MOIs, meropenem treatment did not greatly reduce the intracellular bacterial load nor eradicate all non-persister bacteria. Only at MOIs<1, two or more strains showed no viable bacteria in either the extracellular or intracellular space for at least 2-hours continuously after adding meropenem. Hence, we considered these infected macrophages at MOIs<1 as recovered from infection. As bacterial loads can reflect disease severities [38–43], we classify meropenem-treated, infected macrophages at MOI=1 and MOI=100, as still suffering from mild and severe infections respectively.

### 3.3 Bactericide induced by antibiotics is dependent on bacterial load and differs in the host presence

Aligning to our aim in assessing bactericidal dynamics, we next compared these infection models to either antibiotics alone or macrophages alone for respective strains (**Figure-3**). Bactericidal effects on extracellular bacteria vary for each strain after meropenem treatment in mild infections (**Figure-3A**). At 4-hours post-meropenem, the extracellular bacteria of the EC0210 strain were reduced to from 10^5.21^ to 10^4^ CFU/mL. This magnitude of bacterial reduction is smaller compared to sole meropenem treatment for 4-hours, from 10^5.02^ to 10^1.84^ CFU/mL. A gradual decrease was observed for the EC0238 strain in both meropenem alone (from 10^5.03^ to 10^2^ log CFU/mL) and the extracellular bacteria load of infected macrophages (from 10^4.47^ to 10^1^ log CFU/mL) over the 4-hours duration of meropenem treatment. When the EC0381 strain was treated with solely meropenem, a gradual decrease was observed over the 4-hours of meropenem treatment from 10^5.02^ to 10^2.25^ CFU/mL. Conversely, a rapid decrease of extracellular bacteria from 10^5.56^ to 10^2.46^ CFU/mL was observed at the first hour after adding meropenem, before a bacteriostatic-like effect that ensues. However, no difference was observed between bacteria counts at 4 h when exposed to meropenem and extracellular bacteria counts of infected macrophages at 4 h with exposure to meropenem (*P* = 0.5324) (**Figure-4A**).

**Figure 3:**
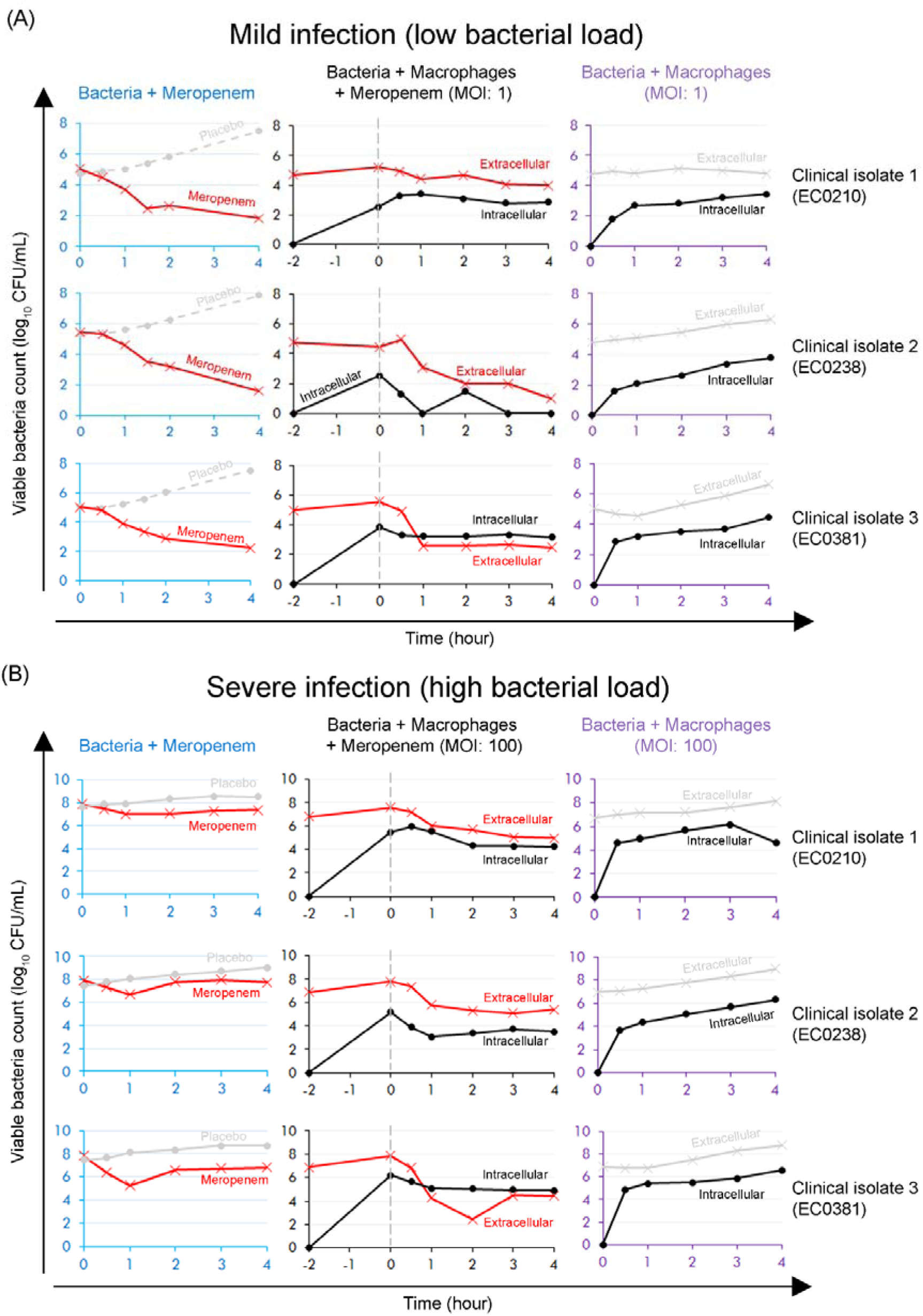
Bactericide by antibiotics is dependent on bacterial load and differs in the host presence. Figure shows models of (A) mild infection (MOI=1) and (B) severe infection (MOI=100). Clinical isolates were exposed to meropenem (20 mg/L) alone, differentiated THP-1 macrophages alone, or meropenem (20 mg/L) and macrophages. Accompanying graphs for side-by-side comparisons shown were either with similar starting inoculums prior to meropenem addition or the same MOIs for infecting THP-1 macrophages. Dotted line represents addition of meropenem after 2 hours post-infection of macrophages by respective clinical strain.

**Figure 4:**
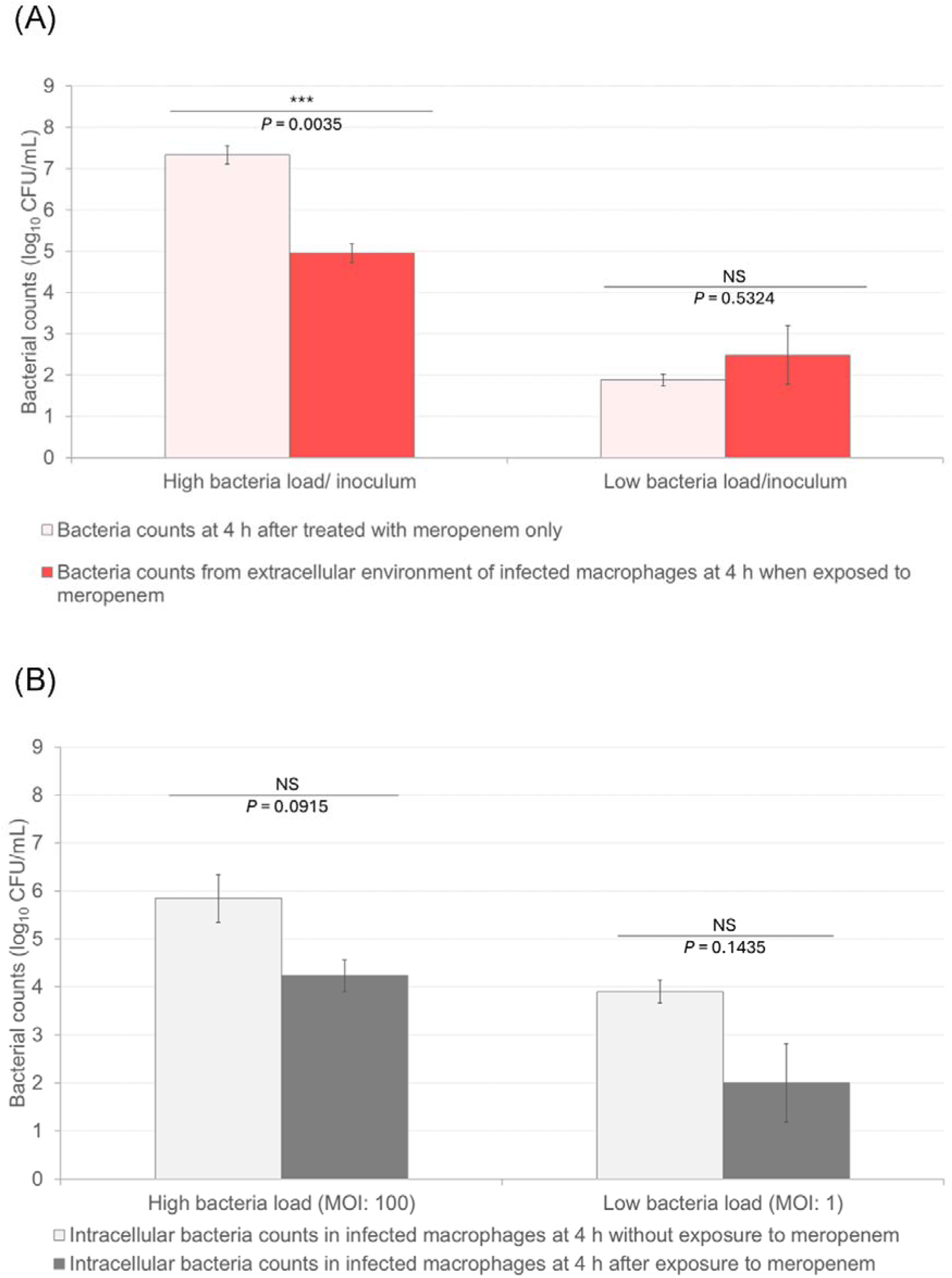
Bacterial counts compared between various treatments at high and low bacteria inoculum/load. (A) Comparisons of bacterial counts at 4 h between meropenem treatment only and extracellular bacteria of macrophage when exposed to meropenem. (B) Comparisons of intracellular bacteria counts in infected macrophages at 4 h between with or without exposure to meropenem. Error bars in this figure represent standard error of the mean. *P*-values were obtained from two-tailed un-paired *t*-test. *** represents *P*-value <0.005; N.S. indicates not statistically significant.

In mild infections (**Figure-3A**), intracellular bacteria numbers reveal bacteriostatic-like effects after 1-hour of meropenem treatment, for both EC0210 and EC0381 isolates. These bacteriostatic-like effects were defined as negligible reductions of bacterial counts with less than 1-logarithmic fold. The intracellular bacteria were reduced from 10^2.52^ CFU/mL to no colony formed for EC0238 strain. Unlike infected macrophages at MOI<1, we do not consider this reduction to zero colony counts as recovered from infection, as the zero colony counts were not maintained for more than one hour.

These intracellular bacteriostatic-like effects contrast against the hyperbolic curves observed with untreated infected macrophages (**Figure-3A**). Complemented with a reduction of extracellular bacteria simultaneously, the bacteriostatic-like effect suggests probable bactericidal events occurring intracellularly. No difference was observed for intracellular bacteria counts in mildly infected macrophages at 4 h when comparing with and without meropenem treatments (*P*=0.1435) (**Figure-4B**). Due to the bacteriostatic-like effect observed intracellularly, we cannot confirm any intracellular, dormant persisters upon meropenem treatment [44,45]. Early studies showed sub-inhibitory meropenem concentrations enhanced macrophage phagocytic activity [46]. As we are unable to quantify non-viable bacteria via colony enumeration, it is inconclusive if meropenem enhances phagocytosis, which includes uptaking of non-viable bacteria or bacterial ghosts killed by meropenem extracellularly.

In severe infections, extracellular viable bacteria of all isolates revealed a sharp decrease with more than 2-logarithmic fold reduction at the first hour of meropenem treatment (**Figure-3B**). A bacteriostatic-like effect then ensues extracellularly. At 4-hours post-meropenem treatment, the extracellular bacteria numbers were consistently 2.4-logarithimic fold lower than sole meropenem exposure across every isolate (*P* = 0.0035), as compared to mild infection which show no significant difference between the same two treatments (*P* = 0.5324) (**Figure-4A**).

In severely infected macrophages (**Figure-3B**), the intracellular bacteria numbers remain relatively constant after 1-2 hours post-meropenem treatment, portraying bacteriostatic-like effects. However, no difference was observed for intracellular bacteria counts in severely infected macrophages at 4 h between with and without meropenem treatments (*P* = 0.0915) (**Figure-4B**).

## 4. DISCUSSION

Using antibiotics for treating bacterial infections is a pivotal moment in modern medicine, transforming the once-lethal infections into treatable conditions and paving the way for safer surgeries, cancer therapies, and organ transplants. Antibiotics are widely recognized to reduce overall bacterial load [1–4], while the remanent bacteria were cleared by the immune system [47].The bacterial load influences both the severity of infection/disease and the effectiveness of immune-mediated clearance.[48,49] Associating the inoculum size/bacterial load to the bacterial load saturation capacity of the host, is essential to understanding not just if an infection will occur, but also how the infection will progress, particularly in the presence of immune or pharmacologic pressures such as antibiotics.

Consistent with studies using cell lines [50] and primary cells [51], our data show bacteria saturation loads in macrophages increases with increasing MOIs (multiplicities of infection, bacteria-to-macrophage ratio). At high MOIs, macrophages can also exhibit reduced killing capacity. Overwhelming bacterial loads tax the cytoskeletal machinery of macrophages, further hindering phagocytosis and intracellular trafficking. This may even trigger stress responses leading to macrophage cell death (e.g. phagocytosis-induced cell death [52])

As previously mentioned, antibiotics were thought to reduce reduces overall bacterial load [1–4], while the remanent bacteria were eliminated by the immune system [47]. Our findings contradict this common notion that antibiotics generally reduce the bacterial load for immune clearance. If the notion holds true for all cases, the bacteria reduction by meropenem should be relatively similar in both the presence and absence of macrophages. Yet, only in severe infections, there were higher bacteria reduction of extracellular bacteria in meropenem treated severe infections compared to bacteria exposed to solely meropenem (**Figure-3B**). Our findings aligned with previous studies showing that synergistic actions of antibiotic and immune cells are context-dependent.[47,53] Antibiotics may affect immune efficiency or trigger complex host-bacteria dynamics that modulate bacterial clearance. Compounding the issue, bacteria can also adopt strategies to resist being destroyed by antibiotics, such as becoming antibiotic tolerant persisters.

Persisters are a sub-population of antibiotic susceptible bacteria that survive antibiotics at bactericidal concentrations. Bactericidal actions by antibiotics is required for identifying persisters *in vitro*, in the absence of host.[14] There are increasing number of recent studies examining host-bacterial persisters interactions of various microbial species [19–27]. A common mechanism of persister formation, in the presence of host, is the bacterial response against reactive nitrogen species from host.[54] Reactive nitrogen species, produced by the host, targeting against the pathogen sequestered within the phagolysosome, can cause bacteria to suppress metabolism, depletes ATP causing bacterial growth arrest and increased antibiotic tolerance.[55–57] Other host-derived stressors, such as nutrient starvation by host (e.g. glucose deprivation) [58] and intraphagosomal environment [54,59] drives persister formation in host.

Our study showing the difference and trend of bacterial reduction between mild and severe infections suggests that antibiotic-induced persisters (not implying intracellular persisters specifically) is formed differently in the presence of the host. We proposed that the formation of antibiotic-induced persisters in host, differs on a case-by-case basis, attributed by infection severity. Physiological change in macrophage (such as being over-whelmed beyond the saturation capacity) may have driven persisters formation differently. This is supported by previous studies demonstrating that different macrophage polarization phenotypes are associated with the presence of either dormant persisters or replicative bacteria [25,60]

Our studies primarily focused on the bacterial infection and clearance dynamics. We did not evaluate the changes in host such as the metabolic reprogramming and surface marker expression, that shroud another layer of complexity in understanding host-persister interactions. One such phenotypic change in host is the formation of macrophage extracellular traps (METs) [61], analogous to neutrophil extracellular traps, that has been reported under high bacterial load. Beyond the classical roles of macrophages in host defence (such as polarizing into the pro-inflammatory M1 phenotype), METs represent a suicidal response that traps pathogens and amplifies inflammation.[61] However, we did not observe morphological features indicative of METs in our study. Instead, we observed differential saturation thresholds across MOIs, suggesting physiological changes in macrophages that warrant further investigation.

In conclusion, bactericidal dynamics of various inoculum sizes differ between the isolates when subjected to both immune and antibiotic pressures, as compared to only exposing to either the antibiotic or the immune cells alone. Contrary to our hypothesis, meropenem did not reduce the intracellular bacterial load within macrophages, regardless of inoculum size. However, meropenem significantly reduced the extracellular bacteria of the macrophages at high inoculum, but not at low inoculum. These findings suggests that antibiotic efficacy in patients may vary from case-to-case, influenced by factors such as infection severity and the specific bacterial isolate causing the infection. We propose that antibiotic-induced persisters (not implying intracellular persisters specifically) formation is different, in the presence of the host, attributed by the infection severity and bacterial load burden.

## DECLARATION

### ETHICS STATEMENT

These bacteria isolates were originally obtained from Singapore General Hospital’s Diagnostic Bacteriology Laboratory, Singapore as part of a surveillance study of carbapenem non-susceptible Gram-negative organisms. No human participants or human participant’s data is involved. Isolates used were de-identified prior to being used in this study.

### DATA AVAILABILITY STATEMENT

All relevant data are within the manuscript and its Supporting Information files.

## ACKNOWLEDGEMENTS

The authors would like to thank the staff of the Division of Pharmacy and Diagnostic Bacteriology Laboratory of the Department of Microbiology at the Singapore General Hospital in their support for this study.

## AUTHOR’S CONTRIBUTION STATEMENT

YJH was involved project design, wrote and edited the manuscript. RLWT and NB were involved for all experimental work required for this study and assist in data analyses. ZY was involved in developing the methodology and optimising the protocol. AKLH was involved in supervision, provided critical input and manuscript review. The final manuscript was read and approved by all contributors.

## Abbreviations (if any)

ATP: Adenosine Triphosphate
CFU: colony forming unit
MET: macrophage extracellular traps
MIC: minimum inhibitory concentrations
MOI: multiplicities-of-infection
PICD: phagocytosis induced cell death

## Supplementary information

**1. Figure S1: Visual validation on uptake of bacteria by THP-1 macrophages**

Representative image of a macrophage infection assay using clinical isolate (EC0210). Bacteria were stained with 150 µM Carboxyfluorescein succinimidyl ester (CFSE) before infecting adhered THP-1 macrophages for 1 h at a MOI of 1. Extracellular cells were washed to remove any unbound bacteria and visually validated under inverted light microscope. Cells were then fixed in 4% (v/v) paraformaldehyde prior for 30 mins, prior to mounting onto glass slides. Images were acquired using a Nikon Widefield microscope. Stained bacteria were visualised using a Nikon-Ti microscope equipped with a Splan Fluor ELWD 40x dry objective lens (Numerical Aperture = 0.6), Nikon Intensilight illumination (excitation) source and a Nikon DS-Ri1-U3 camera. Emitted fluorescence was observed using conventional FITC filters equipped on the microscope at an exposure time of 600 ms. Micrographs acquired were exported as .tif files using the NIS-elements AR software (version 4.2). Corresponding controls with uninfected macrophage and unstain bacteria were also shown. Scale bar represents 10 µm.

## REFERENCES

[1] S. Carryn, F. Van Bambeke, M.P. Mingeot-Leclercq, P.M. Tulkens, Comparative intracellular (THP-1 macrophage) and extracellular activities of β-lactams, azithromycin, gentamicin, and fluoroquinolones against Listeria monocytogenes at clinically relevant concentrations, Antimicrob. Agents Chemother. 46 (2002) 2095–2103. 10.1128/AAC.46.7.2095-2103.2002.

[2] Y. Zhang, Persisters, persistent infections and the Yin-Yang model, Emerg. Microbes Infect. 3 (2014) 1–10. 10.1038/emi.2014.3.

[3] S. Lemaire, F. Van Bambeke, M.P. Mingeot-Leclercq, P.M. Tulkens, Activity of three β-lactams (ertapenem, meropenem and ampicillin) against intraphagocytic Listeria monocytogenes and Staphylococcus aureus, J. Antimicrob. Chemother. 55 (2005) 897–904. 10.1093/jac/dki094.

[4] S. Carryn, F. Van Bambeke, M.-P. Mingeot-Leclercq, P.M. Tulkens, Activity of beta-lactams (ampicillin, meropenem), gentamicin, azithromycin and moxifloxacin against intracellular Listeria monocytogenes in a 24 h THP-1 human macrophage model., J. Antimicrob. Chemother. 51 (2003) 1051–2. 10.1093/jac/dkg189.

[5] S. Helaine, E. Kugelberg, Bacterial persisters: formation, eradication, and experimental systems, Trends Microbiol. 22 (2014) 417–424. 10.1016/j.tim.2014.03.008.

[6] D.M. Monack, A. Mueller, S. Falkow, Persistent bacterial infections: The interface of the pathogen and the host immune system, Nat. Rev. Microbiol. 2 (2004) 747–765. 10.1038/nrmicro955.

[7] R.M. Jones, M. Nicas, A.E. Hubbard, A.L. Reingold, The Infectious Dose of Coxiella Burnetii (Q Fever), Appl. Biosaf. 11 (2006) 32–41. 10.1177/153567600601100106.

[8] R.M. Jones, M. Nicas, A. Hubbard, M.D. Sylvester, A. Reingold, The Infectious Dose of Francisella Tularensis (Tularemia), Appl. Biosaf. 10 (2005) 227–239. 10.1177/153567600501000405.

[9] J. Tuttle, T. Gomez, M.P. Doyle, J.G. Wells, T. Zhao, R. V. Tauxe, P.M. Griffin, Lessons from a large outbreak of Escherichia coli O157:H7 infections: Insights into the infectious dose and method of widespread contamination of hamburger patties, Epidemiol. Infect. 122 (1999) 185–192. 10.1017/S0950268898001976.

[10] H.L. DuPont, M.M. Levine, R.B. Hornick, S.B. Formal, Inoculum Size in Shigellosis and Implications for Expected Mode of Transmission, J. Infect. Dis. 159 (1989) 1126–1128. 10.1093/infdis/159.6.1126.

[11] Y. Hara-Kudo, K. Takatori, Contamination level and ingestion dose of foodborne pathogens associated with infections, Epidemiol. Infect. 139 (2011) 1505–1510. 10.1017/S095026881000292X.

[12] A.B. Kaiser, D.S. Kernodle, R.A. Parker, Low-Inoculum Model of Surgical Wound Infection, J. Infect. Dis. 166 (1992) 393–399. 10.1093/infdis/166.2.393.

[13] A. Brauner, O. Fridman, O. Gefen, N.Q. Balaban, Distinguishing between resistance, tolerance and persistence to antibiotic treatment, Nat. Rev. Microbiol. 14 (2016) 320–330. 10.1038/nrmicro.2016.34.

[14] N.Q. Balaban, S. Helaine, K. Lewis, M. Ackermann, B. Aldridge, D.I. Andersson, M.P. Brynildsen, D. Bumann, A. Camilli, J.J. Collins, C. Dehio, S. Fortune, J.M. Ghigo, W.D. Hardt, A. Harms, M. Heinemann, D.T. Hung, U. Jenal, B.R. Levin, J. Michiels, G. Storz, M.W. Tan, T. Tenson, L. Van Melderen, A. Zinkernagel, Definitions and guidelines for research on antibiotic persistence, Nat. Rev. Microbiol. 17 (2019) 441–448. 10.1038/s41579-019-0196-3.

[15] K. Lewis, Persister cells, dormancy and infectious disease, Nat. Rev. Microbiol. 5 (2007) 48–56. 10.1038/nrmicro1557.

[16] M. Fauvart, V.N. de Groote, J. Michiels, Role of persister cells in chronic infections: Clinical relevance and perspectives on anti-persister therapies, J. Med. Microbiol. 60 (2011) 699–709. 10.1099/jmm.0.030932-0.

[17] P. Bahmaninejad, S. Ghafourian, M. Mahmoudi, A. Maleki, N. Sadeghifard, B. Badakhsh, Persister cells as a possible cause of antibiotic therapy failure in Helicobacter pylori, JGH Open 5 (2021) 493–497. 10.1002/jgh3.12527.

[18] L.R. Mulcahy, J.L. Burns, S. Lory, K. Lewis, Emergence of Pseudomonas aeruginosa strains producing high levels of persister cells in patients with cystic fibrosis, J. Bacteriol. 192 (2010) 6191–6199. 10.1128/JB.01651-09.

[19] A. Xia, A. Thai, Z. Cao, X. Chen, J. Chen, B. Bacacao, L.A. Bekale, V. Schiel, P.L. Bollyky, P.L.S. Maria, Chronic suppurative otitis media causes macrophage-associated sensorineural hearing loss, J. Neuroinflammation 19 (2022) 1–15. 10.1186/s12974-022-02585-w.

[20] M. Hernandez-Morfa, N.M. Reinoso-Vizcaíno, N.B. Olivero, V.E. Zappia, P.R. Cortes, A. Jaime, J. Echenique, Host Cell Oxidative Stress Promotes Intracellular Fluoroquinolone Persisters of Streptococcus pneumoniae, Microbiol. Spectr. 10 (2022). 10.1128/spectrum.04364-22.

[21] J.E. Beam, N.J. Wagner, J.C. Shook, E.S.M. Bahnson, V.G. Fowler, S.E. Rowe, B.P. Conlon, Macrophage-Produced Peroxynitrite Induces Antibiotic Tolerance and Supersedes Intrinsic Mechanisms of Persister Formation, Infect. Immun. 89 (2021). 10.1128/IAI.00286-21.

[22] F. Peyrusson, H. Varet, T.K. Nguyen, R. Legendre, O. Sismeiro, J.Y. Coppée, C. Wolz, T. Tenson, F. Van Bambeke, Intracellular Staphylococcus aureus persisters upon antibiotic exposure, Nat. Commun. 11 (2020). 10.1038/s41467-020-15966-7.

[23] S.E. Rowe, N.J. Wagner, L. Li, J.E. Beam, A.D. Wilkinson, L.C. Radlinski, Q. Zhang, E.A. Miao, B.P. Conlon, Reactive oxygen species induce antibiotic tolerance during systemic Staphylococcus aureus infection, Nat. Microbiol. 5 (2020) 282–290. 10.1038/s41564-019-0627-y.

[24] S. Helaine, A.M. Cheverton, K.G. Watson, L.M. Faure, S.A. Matthews, D.W. Holden, Internalization of salmonella by macrophages induces formation of nonreplicating persisters, Science (80-). 343 (2014) 204–208. 10.1126/science.1244705.

[25] D.A.C. Stapels, P.W.S. Hill, A.J. Westermann, R.A. Fisher, T.L. Thurston, A.E. Saliba, I. Blommestein, J. Vogel, S. Helaine, Salmonella persisters undermine host immune defenses during antibiotic treatment, Science (80-). 362 (2018) 1156–1160. 10.1126/science.aat7148.

[26] N.A. Mayansky, Y.A. Bocharova, E.A. Brzhozovskaya, A. V. Lazareva, I. V. Chebotar, Reactivity of neutrophil-like HL-60 cells towards persistent forms of escherichia coli, Sovrem. Tehnol. v Med. 11 (2019) 82–85. 10.17691/stm2019.11.4.09.

[27] J.M. Mouton, S. Helaine, D.W. Holden, S.L. Sampson, Elucidating population-wide mycobacterial replication dynamics at the single-cell level, Microbiol. (United Kingdom) 162 (2016) 966–978. 10.1099/mic.0.000288.

[28] N.B. Greve, H.C. Slotved, J.E. Olsen, L.E. Thomsen, Identification of antibiotic induced persister cells in Streptococcus agalactiae, PLoS One 19 (2024) 1–13. 10.1371/journal.pone.0303271.

[29] S. Liu, Y. Huang, S. Jensen, P. Laman, G. Kramer, S.A.J. Zaat, S. Brul, S. Liu, Y. Huang, S. Jensen, P. Laman, G. Kramer, S.A.J. Zaat, S. Brul, persister formation in Staphylococcus aureus, (2023).

[30] N. Hofsteenge, E. Van Nimwegen, O.K. Silander, Quantitative analysis of persister fractions suggests different mechanisms of formation among environmental isolates of E. coli, BMC Microbiol. 13 (2013). 10.1186/1471-2180-13-25.

[31] J.H. Yeo, N. Begam, W.T. Leow, J.X. Goh, Y. Zhong, Y. Cai, A.L.-H. Kwa, Ironing out Persisters? Revisiting the Iron Chelation Strategy to Target Planktonic Bacterial Persisters Harboured in Carbapenem-Resistant Escherichia coli., Microorganisms 12 (2024) 972. 10.3390/microorganisms12050972.

[32] T. Starr, T.J. Bauler, P. Malik-Kale, O. Steele-Mortimer, The phorbol 12-myristate-13-acetate differentiation protocol is critical to the interaction of THP-1 macrophages with Salmonella Typhimurium, PLoS One 13 (2018) 1–13. 10.1371/journal.pone.0193601.

[33] D. Petráčková, M.R. Farman, F. Amman, I. Linhartová, A. Dienstbier, D. Kumar, J. Držmíšek, I. Hofacker, M.E. Rodriguez, B. Večerek, Transcriptional profiling of human macrophages during infection with Bordetella pertussis, RNA Biol. 17 (2020) 731–742. 10.1080/15476286.2020.1727694.

[34] V.H. Tam, A.N. Schilling, M. Nikolaou, Modelling time-kill studies to discern the pharmacodynamics of meropenem, J. Antimicrob. Chemother. 55 (2005) 699–706. 10.1093/jac/dki086.

[35] CLSI, Performance Standards for Antimicrobial Susceptibility Testing. 30th Edition. CLSI supplement M100., 30th ed., Clinical and Laboratory Standards Institute, Wayne, PA 19087, USA, 2020.

[36] I.M. Ghazi, J.L. Crandon, E.P. Lesho, P. McGann, D.P. Nicolau, Efficacy of Humanized High-Dose Meropenem, Cefepime, and Levofloxacin against Enterobacteriaceae Isolates Producing Verona Integron-Encoded Metallo-β-Lactamase (VIM) in a Murine Thigh Infection Model, Antimicrob. Agents Chemother. 59 (2015) 7145–7147. 10.1128/AAC.00794-15.

[37] F.S. Taccone, F. Cotton, S. Roisin, J.L. Vincent, F. Jacobs, Optimal meropenem concentrations to treat multidrug-resistant Pseudomonas aeruginosa septic shock, Antimicrob. Agents Chemother. 56 (2012) 2129–2131. 10.1128/AAC.06389-11.

[38] I. Stranieri, K.A. Kanunfre, J.C. Rodrigues, L. Yamamoto, M.I.V. Nadaf, P. Palmeira, T.S. Okay, Assessment and comparison of bacterial load levels determined by quantitative amplifications in blood culture-positive and negative neonatal sepsis, Rev. Inst. Med. Trop. Sao Paulo 60 (2018) e61. 10.1590/S1678-9946201860061.

[39] O. Sibila, E. Laserna, A. Shoemark, H.R. Keir, S. Finch, A. Rodrigo-Troyano, L. Perea, M. Lonergan, P.C. Goeminne, J.D. Chalmers, Airway bacterial load and inhaled antibiotic response in bronchiectasis, Am. J. Respir. Crit. Care Med. 200 (2019) 33–41. 10.1164/rccm.201809-1651OC.

[40] N.T.T. Thuong, D.N. Vinh, H.T. Hai, D.D.A. Thu, L.T.H. Nhat, D. Heemskerk, N.D. Bang, M. Caws, N.T.H. Mai, G.E. Thwaites, Pretreatment cerebrospinal fluid bacterial load correlates with inflammatory response and predicts neurological events during tuberculous meningitis treatment, J. Infect. Dis. 219 (2019) 986–995. 10.1093/infdis/jiy588.

[41] M.R. Spalinger, V. Canale, A. Becerra, A. Shawki, M. Crawford, A.N. Santos, P. Chatterjee, J. Li, M.G. Nair, D.F. McCole, PTPN2 regulates bacterial clearance in a mouse model of enteropathogenic and enterohemorrhagic E. coli infection, JCI Insight 8 (2023). 10.1172/jci.insight.156909.

[42] A. Kumar, D. Roberts, K.E. Wood, B. Light, J.E. Parrillo, S. Sharma, R. Suppes, D. Feinstein, S. Zanotti, L. Taiberg, D. Gurka, A. Kumar, M. Cheang, Duration of hypotension before initiation of effective antimicrobial therapy is the critical determinant of survival in human septic shock, Crit. Care Med. 34 (2006) 1589–1596. 10.1097/01.CCM.0000217961.75225.E9.

[43] M. Varbanova, P. Malfertheiner, Bacterial load and degree of gastric mucosal inflammation in helicobacter pylori infection, Dig. Dis. 29 (2011) 592–599. 10.1159/000333260.

[44] N.J. Bokil, M. Totsika, A.J. Carey, K.J. Stacey, V. Hancock, B.M. Saunders, T. Ravasi, G.C. Ulett, M.A. Schembri, M.J. Sweet, Intramacrophage survival of uropathogenic Escherichia coli: Differences between diverse clinical isolates and between mouse and human macrophages, Immunobiology 216 (2011) 1164–1171. 10.1016/j.imbio.2011.05.011.

[45] K. Sharma, N. Dhar, V. V. Thacker, T.M. Simonet, F. Signorino-Gelo, G. Knott, J.D. McKinney, Dynamic persistence of intracellular bacterial communities of uropathogenic escherichia coli in a human bladder-chip model of urinary tract infections, Elife 10 (2021) 1–30. 10.7554/eLife.66481.

[46] A.M. Cuffini, V. Tullio, A. Allocco, F. Giachino, S. Fazari, N.A. Carlone, The entry of meropenem into human macrophages and its immunomodulating activity, J. Antimicrob. Chemother. 32 (1993) 695–703. 10.1093/jac/32.5.695.

[47] P. Ankomah, B.R. Levin, Exploring the collaboration between antibiotics and the immune response in the treatment of acute, self-limiting infections, Proc. Natl. Acad. Sci. U. S. A. 111 (2014) 8331–8338. 10.1073/pnas.1400352111.

[48] D. Pessoa, C. Souto-Maior, E. Gjini, J.S. Lopes, B. Ceña, C.T. Codeço, M.G.M. Gomes, Unveiling Time in Dose-Response Models to Infer Host Susceptibility to Pathogens, PLoS Comput. Biol. 10 (2014). 10.1371/journal.pcbi.1003773.

[49] L. Ralberg, D. Sim, A.F. Read, Disentangling Genetic Variation for Resistance and Tolerance to Infectious Diseases in Animals, Science (80-). 318 (2007) 812–814. 10.1126/science.1148526.

[50] M.J. Olivares-Morales, M.K. De La Fuente, K. Dubois-Camacho, D. Parada, D. Diaz-Jiménez, A. Torres-Riquelme, X. Xu, N. Chamorro-Veloso, R. Naves, M.J. Gonzalez, R. Quera, C. Figueroa, J.A. Cidlowski, R.M. Vidal, M.A. Hermoso, Glucocorticoids impair phagocytosis and inflammatory response against Crohn’s disease-associated adherent-invasive escherichia coli, Front. Immunol. 9 (2018) 1–14. 10.3389/fimmu.2018.01026.

[51] C. Neu, A. Sedlag, C. Bayer, S. Förster, P. Crauwels, J.H. Niess, G. van Zandbergen, G. Frascaroli, C.U. Riedel, CD14-Dependent Monocyte Isolation Enhances Phagocytosis of Listeria monocytogenes by Proinflammatory, GM-CSF-Derived Macrophages, PLoS One 8 (2013). 10.1371/journal.pone.0066898.

[52] D. El Kebir, J.G. Filep, Modulation of neutrophil apoptosis and the resolution of inflammation through β2 integrins, Front. Immunol. 4 (2013) 1–15. 10.3389/fimmu.2013.00060.

[53] S.M. Lehar, T. Pillow, M. Xu, L. Staben, K.K. Kajihara, R. Vandlen, L. DePalatis, H. Raab, W.L. Hazenbos, J. Hiroshi Morisaki, J. Kim, S. Park, M. Darwish, B.C. Lee, H. Hernandez, K.M. Loyet, P. Lupardus, R. Fong, D. Yan, C. Chalouni, E. Luis, Y. Khalfin, E. Plise, J. Cheong, J.P. Lyssikatos, M. Strandh, K. Koefoed, P.S. Andersen, J.A. Flygare, M. Wah Tan, E.J. Brown, S. Mariathasan, Novel antibody-antibiotic conjugate eliminates intracellular S. aureus, Nature 527 (2015) 323–328. 10.1038/nature16057.

[54] S. Helaine, B.P. Conlon, K.M. Davis, D.G. Russell, Host stress drives tolerance and persistence: The bane of anti-microbial therapeutics, Cell Host Microbe 32 (2024) 852–862. 10.1016/j.chom.2024.04.019.

[55] J. Bayer, J. Becker, X. Liu, L. Gritsch, E. Daiber, N. Korn, F. Oesterhelt, M. Fraunholz, A. Weber, C. Wolz, Differential survival of Staphylococcal species in macrophages, Mol. Microbiol. 121 (2024) 470–480. 10.1111/mmi.15184.

[56] S. Ronneau, C. Michaux, S. Helaine, Decline in nitrosative stress drives antibiotic persister regrowth during infection, Cell Host Microbe 31 (2023) 993–1006.e6. 10.1016/j.chom.2023.05.002.

[57] S. Ronneau, C. Michaux, R.T. Giorgio, S. Helaine, Intoxication of antibiotic persisters by host RNS inactivates their efflux machinery during infection, PLoS Pathog. 20 (2024) 1–16. 10.1371/journal.ppat.1012033.

[58] M. Huemer, S.M. Shambat, J. Bergada-Pijuan, S. Söderholm, M. Boumasmoud, C. Vulin, A. Gómez-Mejia, M.A. Varela, V. Tripathi, S. Götschi, E.M. Maggio, B. Hasse, S.D. Brugger, D. Bumann, R.A. Schuepbach, A.S. Zinkernagel, Molecular reprogramming and phenotype switching in Staphylococcus aureus lead to high antibiotic persistence and affect therapy success, Proc. Natl. Acad. Sci. U. S. A. 118 (2021) 1–12. 10.1073/pnas.2014920118.

[59] I. Dadole, D. Blaha, N. Personnic, The macrophage–bacterium mismatch in persister formation, Trends Microbiol. xx (2024) 1–13. 10.1016/j.tim.2024.02.009.

[60] A.E. Saliba, L. Li, A.J. Westermann, S. Appenzeller, D.A.C. Stapels, L.N. Schulte, S. Helaine, J. Vogel, Single-cell RNA-seq ties macrophage polarization to growth rate of intracellular Salmonella, Nat. Microbiol. 2 (2016) 1–8. 10.1038/nmicrobiol.2016.206.

[61] A.A. Baz, H. Hao, S. Lan, Z. Li, S. Liu, X. Jin, S. Chen, Y. Chu, Emerging insights into macrophage extracellular traps in bacterial infections, FASEB J. 38 (2024) 1–14. 10.1096/fj.202400739R.

