## Supplementary figures and images for "Assessing bactericidal dynamics of persister-manifested clinical isolates in the presence of macrophages and antibiotics"

### Supplementary Figure 1

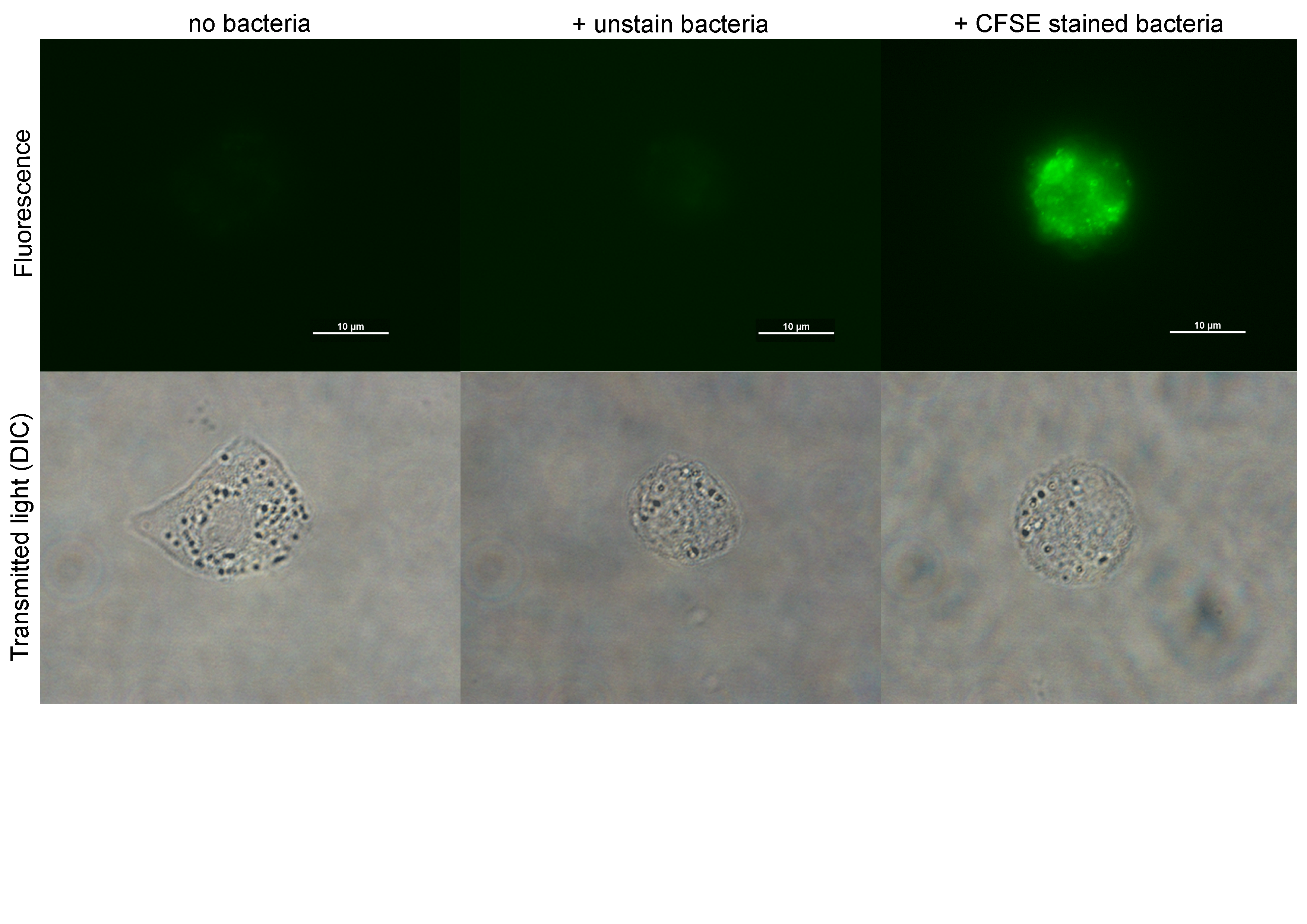
